# The QNOAEL vs. BMD for Point of Departure

**DOI:** 10.1101/329763

**Authors:** Cynthia Dickerson, Robert A. Lodder

## Abstract

Quantile bootstrap (QB) methods can be applied to the problem of estimating the No Observed Adverse Effect Level (NOAEL) of a New Molecular Entity (NME) to anticipate a safe starting dose for beginning clinical trials. An estimate of the NOAEL from the extended QB method (called the QNOAEL) can be calculated using multiple disparate studies in the literature and/or from laboratory experiments. The QNOAEL is similar in some ways to the Benchmark Dose (BMD) and is superior to the BMD in others. The Benchmark Dose method is currently widely used in toxicological research.

Results are used in a simulation based on nonparametric cluster analysis methods to calculate confidence levels on the difference between the Effect and the No Effect studies. The QNOAEL simulation generates an intuitive curve that is comparable to the dose-response curve.

The QNOAEL of ellagic acid (EA) will be calculated for clinical trials of its use as a component therapeutic agent (in BSN476) for treating Chikungunya infections. This will be the first application of QB to the problem of NOAEL estimation for a drug. The specific aims of the proposed study are to evaluate the accuracy and precision of the QB Simulation and QNOAEL compared to the Benchmark Dose Method, and to calculate the QNOAEL of EA for BSN476 Drug Development.

## Specific Aims

Nonparametric statistics are statistics that are <u>not</u> based on parameterized families of probability distributions, like the normal distribution. They are important because data frequently follow a distribution other than a known one, like the normal distribution.

The NOAEL is an important part of the non-clinical risk assessment for new drugs like BSN476, a drug for treating Chikungunya. The NOAEL is a professional opinion based on the design of the study, indication of the drug, expected pharmacology, and spectrum of off-target effects. It is the highest dose at which there was not an observed toxic or adverse effect^1^. There are important theoretical limitations to the traditional NOAEL calculation, which led to the newer Benchmark Dose method, which also has a number of problems. In brief, the traditional NOAEL is determined by administering a few different doses of drug to a group of subjects, observing those subjects for physiological change, and assigning the dosages to the categories of “having an adverse effect” and “not having an adverse effect”. The highest dosage resulting in no adverse effect is determined to be the NOAEL.

The NOAEL method is problematic because (1) dose levels are often an order of magnitude apart, and it is highly unlikely that the exact NOAEL dosage will be administered in any particular study; (2) determination of what constitutes an “effect” can be difficult when negative effects are of a highly subjective nature (for example, when mood is affected)^2^. A nonparametric simulation using extended QB (Quantile Bootstrap) methods can solve the problems associated with the use of the traditional NOAEL or the Benchmark Dose (BMD) and enable accurate toxicological estimates to be made.

### Evaluate the accuracy and precision of the QB Simulation and QNOAEL compared to the Benchmark Dose Method

Utilizing synthetic data with known characteristics, the BMD and QNOAEL will be calculated. The QNOAEL will then be compared to the BMD to determine which is closest to be known answer for the synthetic data.

### Calculate the QNOAEL of Ellagic Acid for BSN476 Drug Development

Chikungunya is a rapidly spreading mosquito-borne disease that now infects over 3 million people worldwide ^3^. BSN476, a drug for treating chikungunya infections, contains in part EA. QB will estimate a safe level of EA for the first-in-human study in order to develop a treatment for Chikungunya. An EA toxicity meta-analysis using food consumption will be completed as part of the Investigational New Drug (IND) application to the FDA. Studies will be selected from the literature and analyzed according to the Cochrane protocols, and the QNOAEL of EA will be calculated along with the NOAEL and BMD. These results will serve as the basis for the first-in-human study of EA.

## Strategy

### Significance

#### Method Significance

The NOAEL depends strongly on the dose selection, dose spacing, and sample size of a single study from which the critical effect has been identified. The primary goal of BMD modeling is to define a point of departure that is largely independent of study design. But while the BMD effectively enables multiple studies to be pooled to increase accuracy, it does not handle studies with conflicting results gracefully, as will be seen below ^4^.

|  | NOAEL | Benchmark Dose | QNOAEL |
| --- | --- | --- | --- |
| Advantages | Very simple, early method of deriving a POD that has been utilized for years<br>Defined as the highest dose which | Calculation is not limited to specific experimental doses, but can calculate a BMD within the tested range<br>Multiple studies may be pooled to increase accuracy<br>Allows for weighting of | The advantages of BMD, plus:<br>Calculates statistical measures of certainty to gracefully handle conflicting input study results<br>Generates an intuitive correlation vs. dose curve that shows the statistical likelihood of effect at a given dose<br>Able to account for publication bias in the literature |
|  | generates no effect in the studied population<br>Can be used when data will not work with BMD model | results based on study quality<br>Accounts for dose-response curve and pharmacokinetic modeling<br>Benchmark Dose is defined as the dose which generates a specific benchmark effect | Bootstrapping algorithm is robust and nonparametric (and thus able to solve problems in which the underlying distribution is unknown )<br>Can extrapolate to lower doses than dosing range studied/reported in the literature.<br>Computed quickly(an order of $n^1$ algorithm) |
| Limitations | Limited to doses studied/reported in the literature<br>Lowest Observed Average Effect Level (LOAEL) cannot be used to generate NOAEL<br>Not designed for pooling of multiple studies<br>Does not take into account the statistical likelihood of effect across a large population<br>Does not represent the population as a whole; is limited to the population studied | More complicated and time-consuming process than simple NOAEL<br>Limited strongly by the quality of information and the inclusion criteria chosen by the researcher<br>Unable to account well for conflicting study results<br>Generates a non-intuitive mean response x dose line which is ambiguous for doses which sometimes cause an effect, or which cause an effect in a certain percentage of the population<br>Results do not well indicate the response of a large population to a drug<br>Parametric test.<br>"Effect" vs. "No Effect" classification can be difficult when studies examined different endpoints; ex: effect on heart rate vs. mood alteration | More complicated and time-consuming process than simple NOAEL<br>Limited somewhat by the quality of information and the study inclusion criteria chosen by the researcher |

The BMDL (the statistical lower confidence limit on the benchmark dose, or BMD) is used as the point of departure (POD) for most non-cancer and cancer risk estimates derived by the U.S. EPA. The initial step in the risk assessment process is hazard identification, which is defined as the identification of effects on health noted as the result of exposure to a specified chemical. Hazard identification is followed by determination of the critical effect on which to construct NOAELs (No Adverse Effect Levels) or BMDs and BMDLs.

Both the NOAEL and BMD approach require some common considerations of the general quality of a particular study. A few of these considerations include:

a. Sample Size. Were the sample sizes used large enough to properly detect treatment effects?
b. Exposure. Were the exposure durations adequate? Were relevant routes of exposure employed in the study?
c. Endpoints. Did the study measure endpoints of interest?
d. Quality. Did the study employ standard quality control procedures like good laboratory practice (GLP)?

In addition to these common data quality considerations that affect both the NOAEL and BMD estimates, there are added BMD-specific points to consider in the identification of datasets that are appropriate for BMD modeling. For example, when sample size decreases, which results in decreased power to detect treatment effects, the NOAEL procedure produces higher POD estimates while the BMD approach produces lower (extra precautionary) PODs. To maintain consistency and reproducibility, most scientists employ a six-step process for BMD analysis. The six steps involved in the BMD analysis are (1) choice of a BMR, (2) selecting a set of models, (3) assessing model fit, (4) model selection when BMDLs are divergent, (5) model selection when BMDLs are not divergent, and (6) data reporting.

The new QB nonparametric meta-analysis of multiple studies so far appears to be superior to BMD modeling. Unlike BMD, the QNOAEL estimate is not limited by the format of the data presented. The QNOAEL is no more time-consuming to calculate than the BMD, and provides a simpler decision-making process. For example, the graphs below show the BMD and QB simulations for THC in hemp seed. Note that the same clinical studies were used for both analyses. QB not only more clearly demonstrates the trend of data, but also produces a correlation curve which is intuitively noncontradictory. (It is to be expected that different studies may reach contradictory results on the effects of a given dosage, as study methods and populations vary.)

**Fig. 1.**
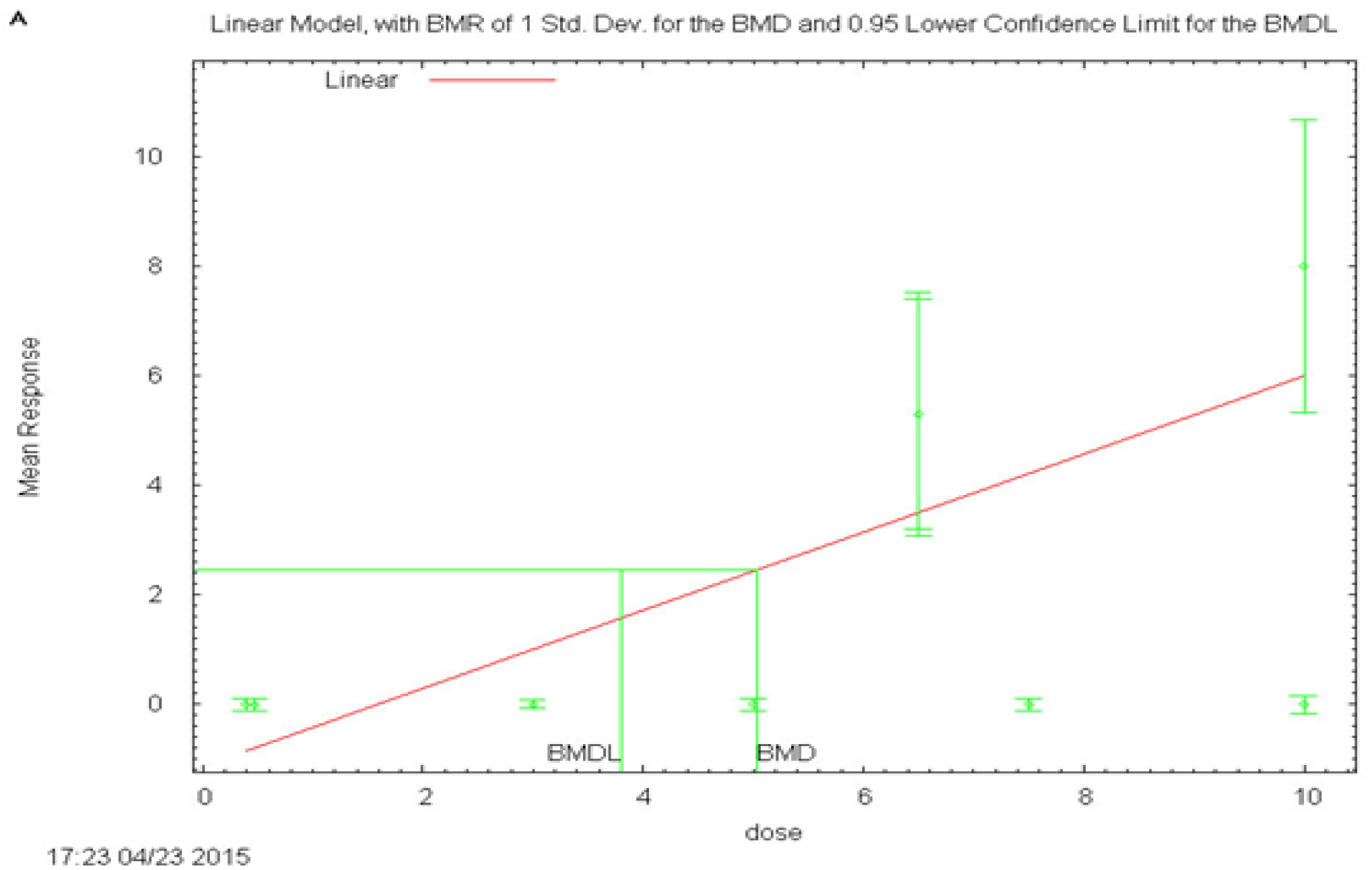
The BMD method attempts to fit a model to studies at different THC doses (green circles) that sometimes show an effect, and sometimes do not. This fitting process can seem to make little sense considering the data. Note that two different studies at 10 mg THC show opposite effects. (Dosages falling on the 0 line of “Mean Response” are no-effect dosages in this figure.)

**Fig. 2.**
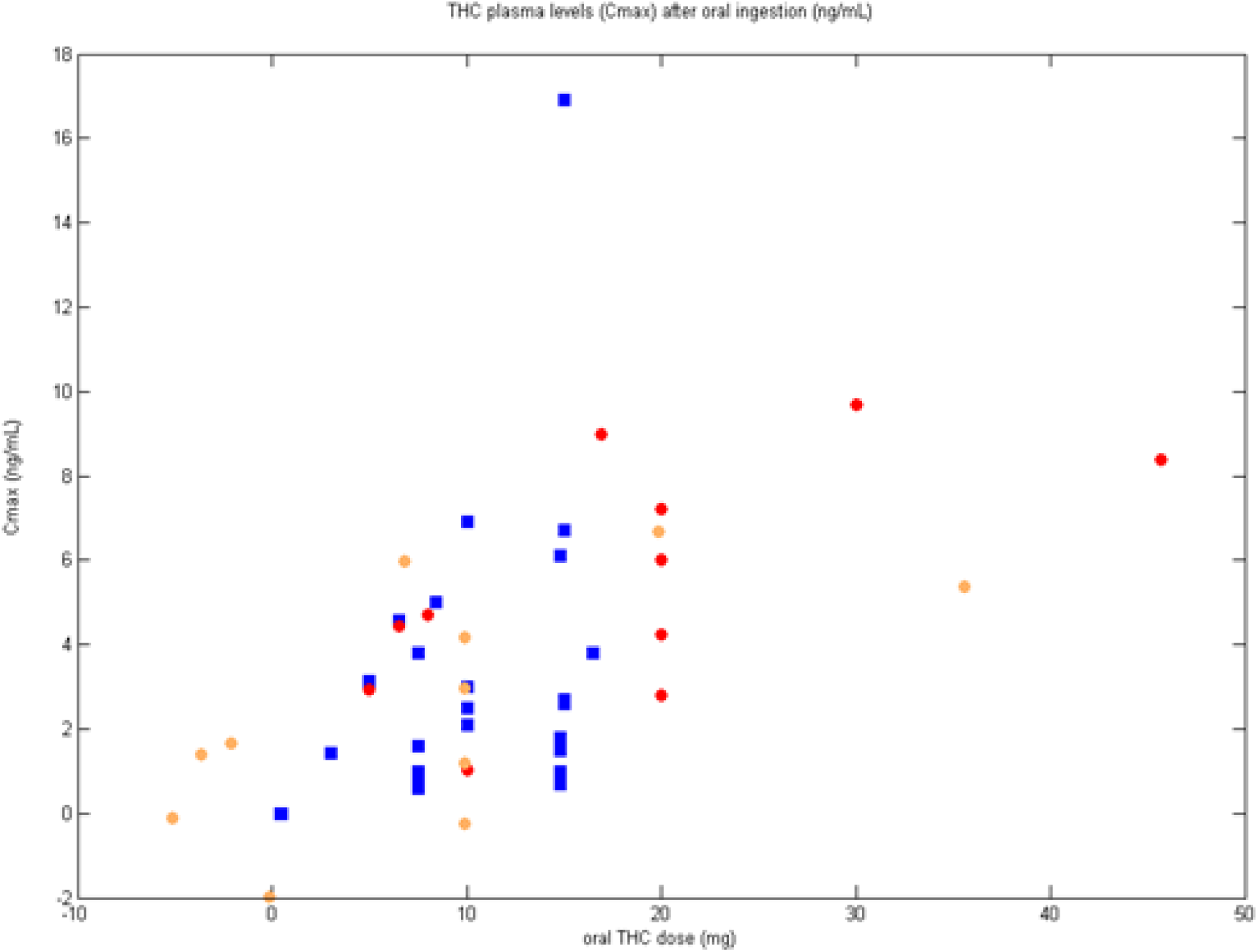
On the other hand, in multivariate space the doses show no discontinuities. Red circles are test set study results (showed effect on heart rate HR and blood pressure BP). Blue squares are training set study results (no effect on HR or BP, sometimes an effect on C_max_). Orange circles are test set after recentering on the training set.

**Fig. 3.**
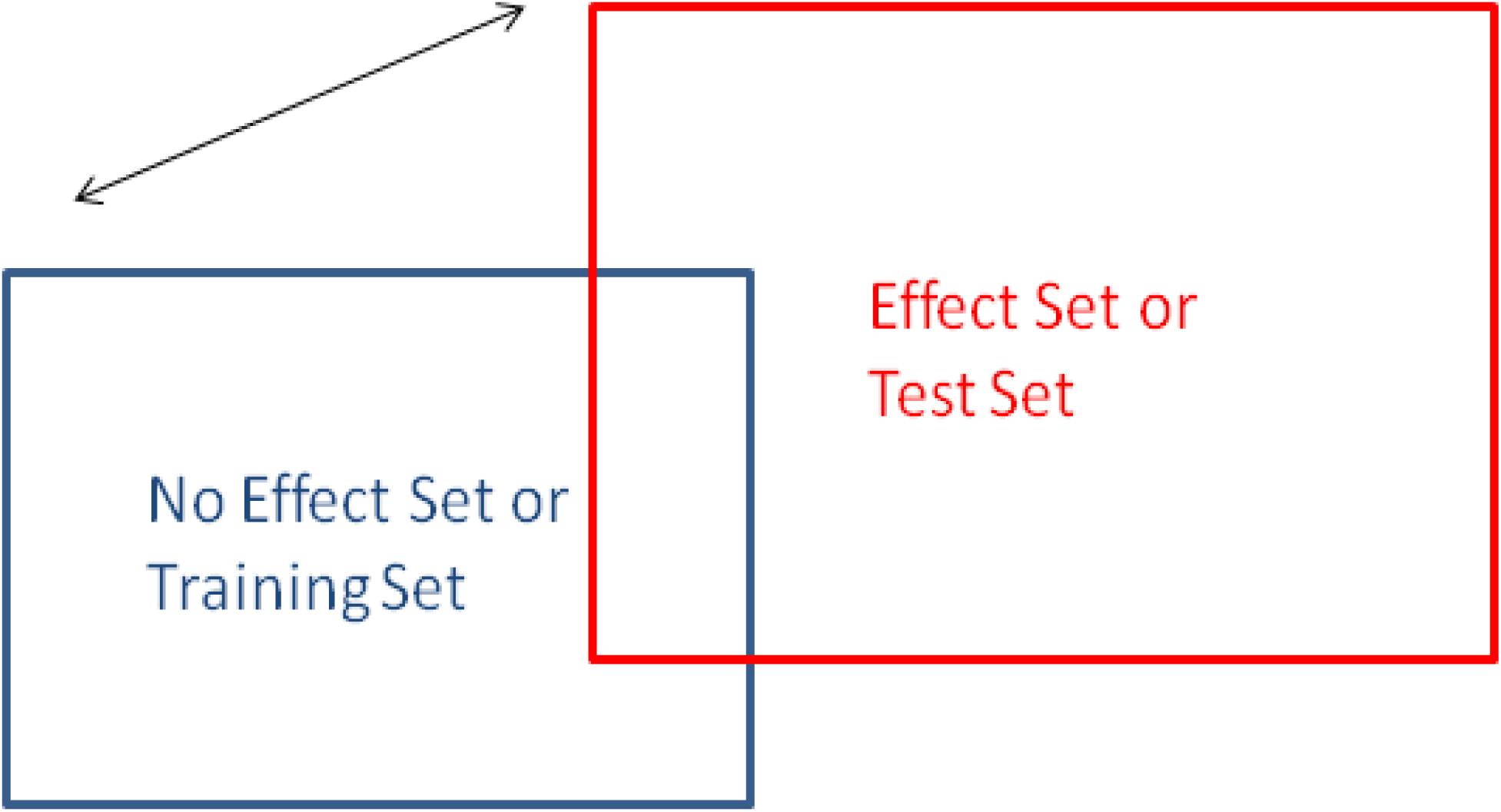
In the new nonparametric method, 98% confidence limits are set on the No Effect Set (or Training Set) by correlating integrals of bootstrap replicates of the training set with themselves. The simulation then projects the Effect Set (or Test Set) into the same space as the No Effect Set, and then translates the Effect Set toward/away from the center of the No Effect Set to determine when the correlations between the No Effect and Effect replicate integrals become significantly different.

**Fig. 4.**
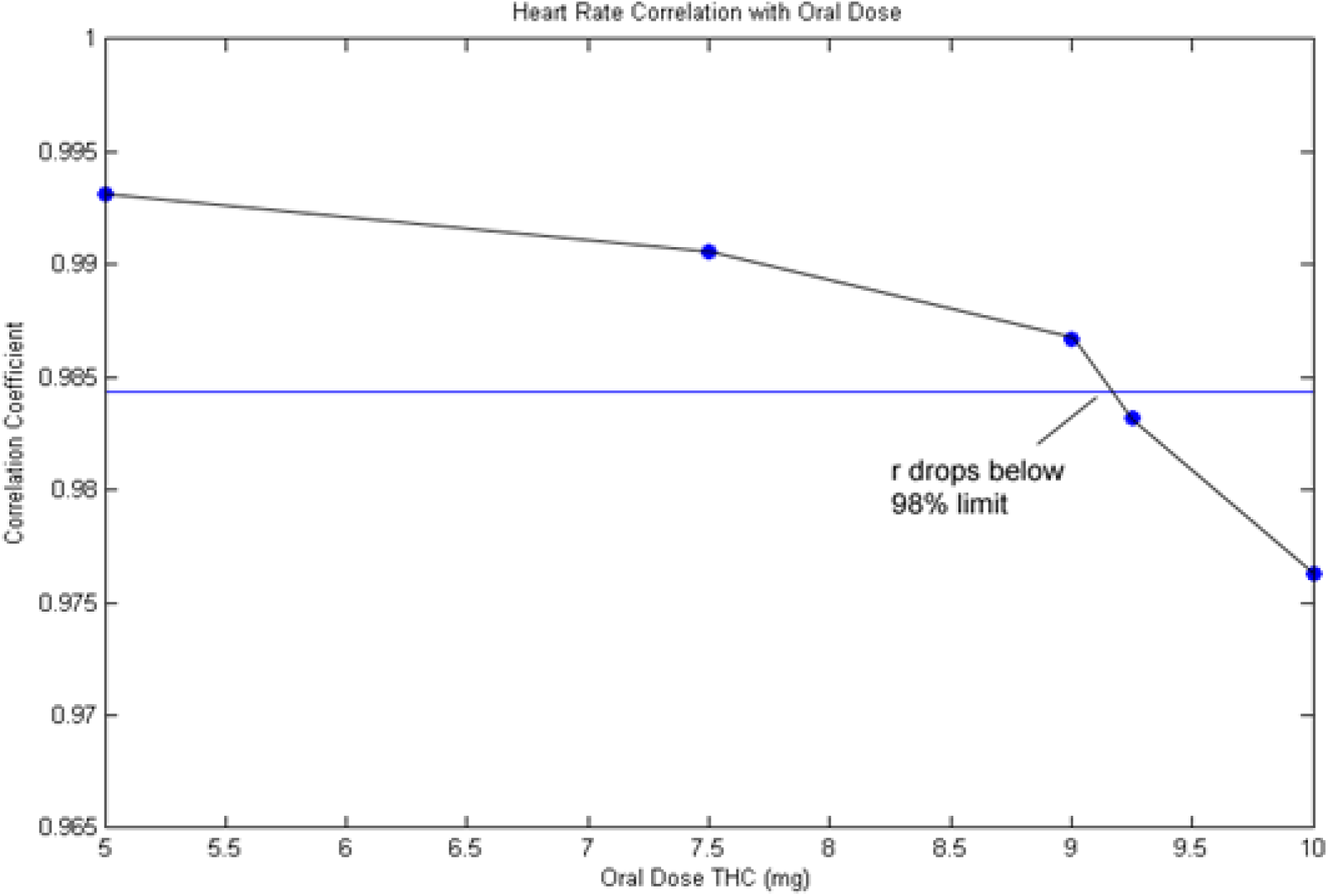
The gradual approach of the line to the 98% confidence limit reveals that the scale of the test set is larger than the scale of the training set. The horizontal blue line represents the 98% confidence limit on the training set (the no effect on HR/BP set). The 98% confidence limit on the training set was 9.1 mg orally (which is the NOEL for heart rate) with a C_max_ = 2.8 ng/ml.

#### Application Significance

Chikungunya is a rapidly spreading mosquito-borne disease that now infects over 3 million people worldwide ^3^. The disease originated in Africa around 1700 A.D., and until recent years, reported infections were limited to the African continent and Southeast Asia ^5^. The disease was first identified in 1952 during an outbreak so serious that infections were clinically indistinguishable from dengue fever ^6^. Throughout the 1960s and 1970’s, outbreaks were reported in Southeast Asia^6^. After decades without another Southeast Asian outbreak, a 1999 outbreak in Indonesia led to a massive outbreak reported in India in 2006, the strain responsible for this resurgence bearing 99% similarity to the strain responsible for a 1989 outbreak in Uganda ^4,6^. In December, 2013, the disease made its debut in the Americas, with its first local transmission occurring on the island of St. Martin; local transmission in French Guiana on the South American continent occurred later that month ^7-9^. After only two years, local transmission had been documented in 19 Caribbean countries, including Puerto Rico, as well as in nearly every country on the South American continent ^7-10^. The WHO has currently issued a level-1 watch for travelers visiting South America and the Caribbean and expects Chikungunya to spread ^7,8^.

BSN476 contains in part EA. EA is a polyphenolic compound with antiproliferative and antiviral properties^11^. EA at 10 uM produces 99.6% inhibition of Chikungunya virus in vitro ^11^. EA is found in a number of plant extracts, usually in the form of hydrolyzable ellagitannins which are complex esters of EA with glucose ^12,13^. Ellagitannins are broken down in the intestine to eventually release EA ^12,13^. To develop BSN476 as a treatment for Chikungunya, the PK of the drug must be studied in a first-in-human (FIH) trial, and a safe range of exposure must be determined for that trial.

**Fig. 5.**
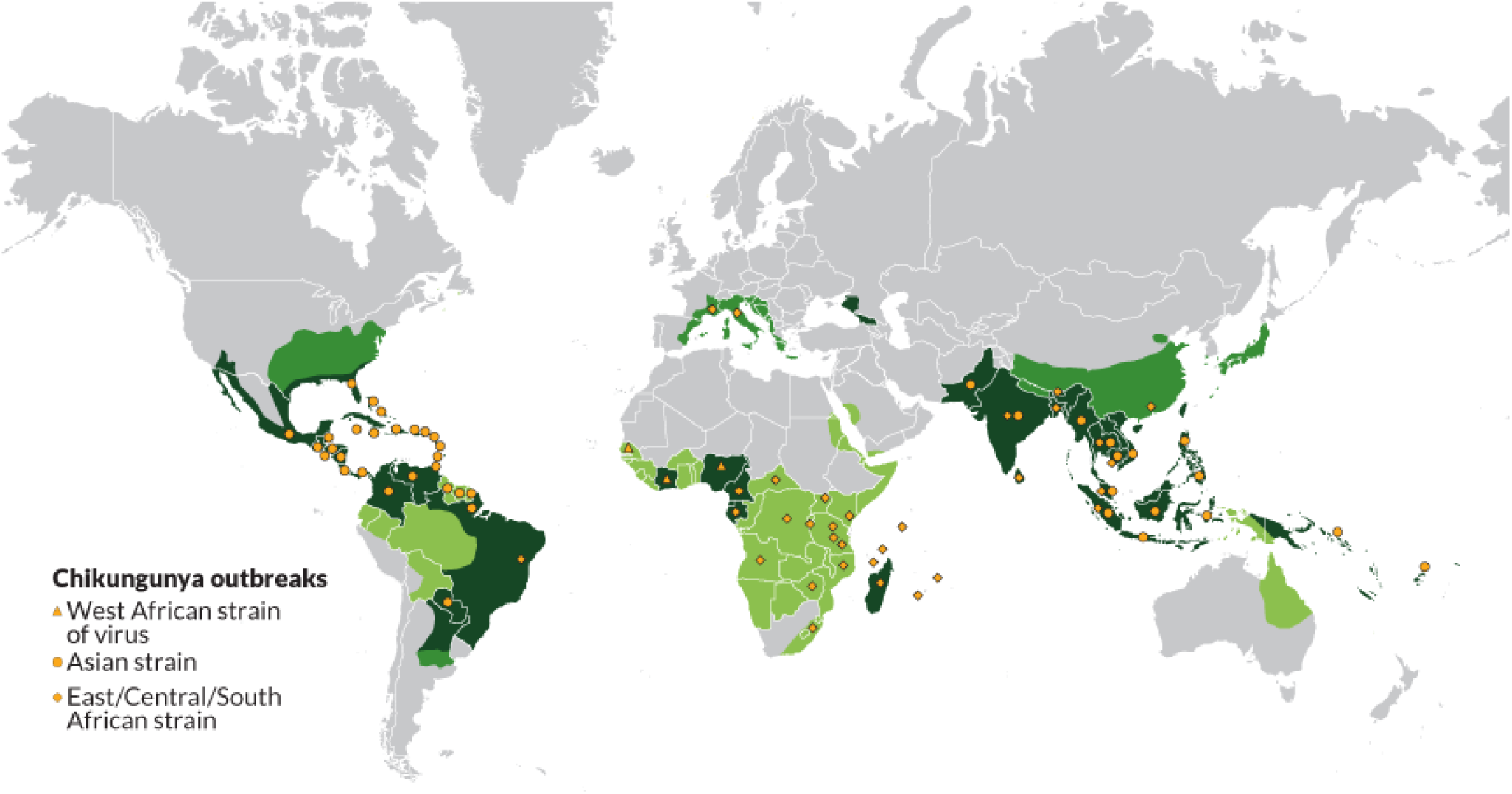
Yellow dots mark the locations of chikungunya outbreaks, while the green areas mark the ranges of the mosquitoes able to propagate the outbreaks. This map was shows data only from June, 2015 and earlier.3

QB is applied within the Cochrane framework for meta-analysis (the Cochrane framework provides a “Garbage-In-Garbage-Out” standard for data inputs - generally clinical studies) to determine doses for the first in human study ^14^. The QNOAEL of EA will be estimated from previously published food consumption studies. Bootstrap replication and manipulation of data clusters will reveal the QNOAEL of EA with 98% confidence.

This project will use the QNOAEL to estimate a safe range of exposure to EA for the FIH study on the way to developing a treatment for Chikungunya. QB is a robust O(n^1^) algorithm that is designed for massively parallel computers, and is a very powerful meta-analysis tool (most algorithms use matrix factorization and are O(n3) in execution time). The QB algorithm will be translated from MATLAB into Python to make it more accessible to the scientific community. Very large sample datasets will be used to test the memory usage and reproducibility of the results of the algorithm, as well as minimum parameters for its usage.

These data have permitted estimation of the maximum analysis capability of cloud computing services and the National Science Foundation XSEDE supercomputer (Comet).

### Innovation

This proposed study utilizes a Quantile Bootstrap statistical method designed for Big Data problems, SOB, inside a new simulation to estimate the QNOAEL for a drug with 98% confidence from a set of small studies ^15,15-18^. SOB is a form of cluster analysis, which is a common analytical technique for determining chemical identity and purity ^15-18^. So far, the QNOAEL appears to be superior to the NOAEL and the BMD.

QB is applied within the Cochrane framework for meta-analysis. QB works by analyzing clusters of studies that found an effect, and clusters of studies that found no effect (the studies can use different dose levels). By analyzing the quantiles of each cluster and adjusting for cluster skew, QB can measure the distance between clusters in probability space, until it finds the dose that yields no adverse effects for the entire human population at a specified level of statistical significance (see Fig. 6, 98% level set by default).

**Fig. 6.**
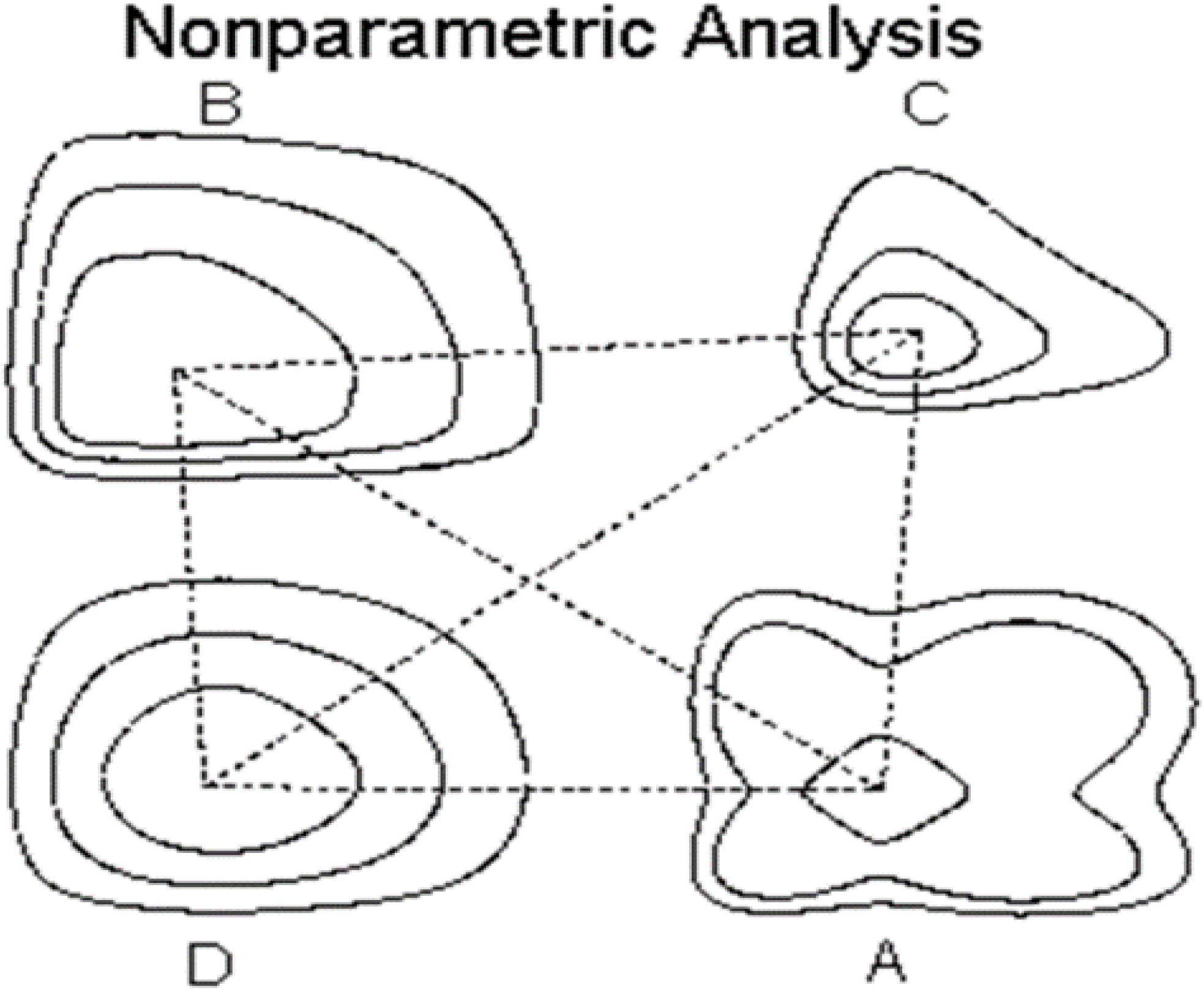
Distances in BEST SDs depend not only on the direction in space, but also on the cluster selected to develop the metric.

### Approach

This is a software development project undertaken as part of a larger drug development project. Good Engineering Practice, Standard Operating Procedures (SOPs) and working practice guidelines have been implemented for project design as well as execution (see Fig. 7). A robust change control system must be implemented in this project.

**Fig. 7.**
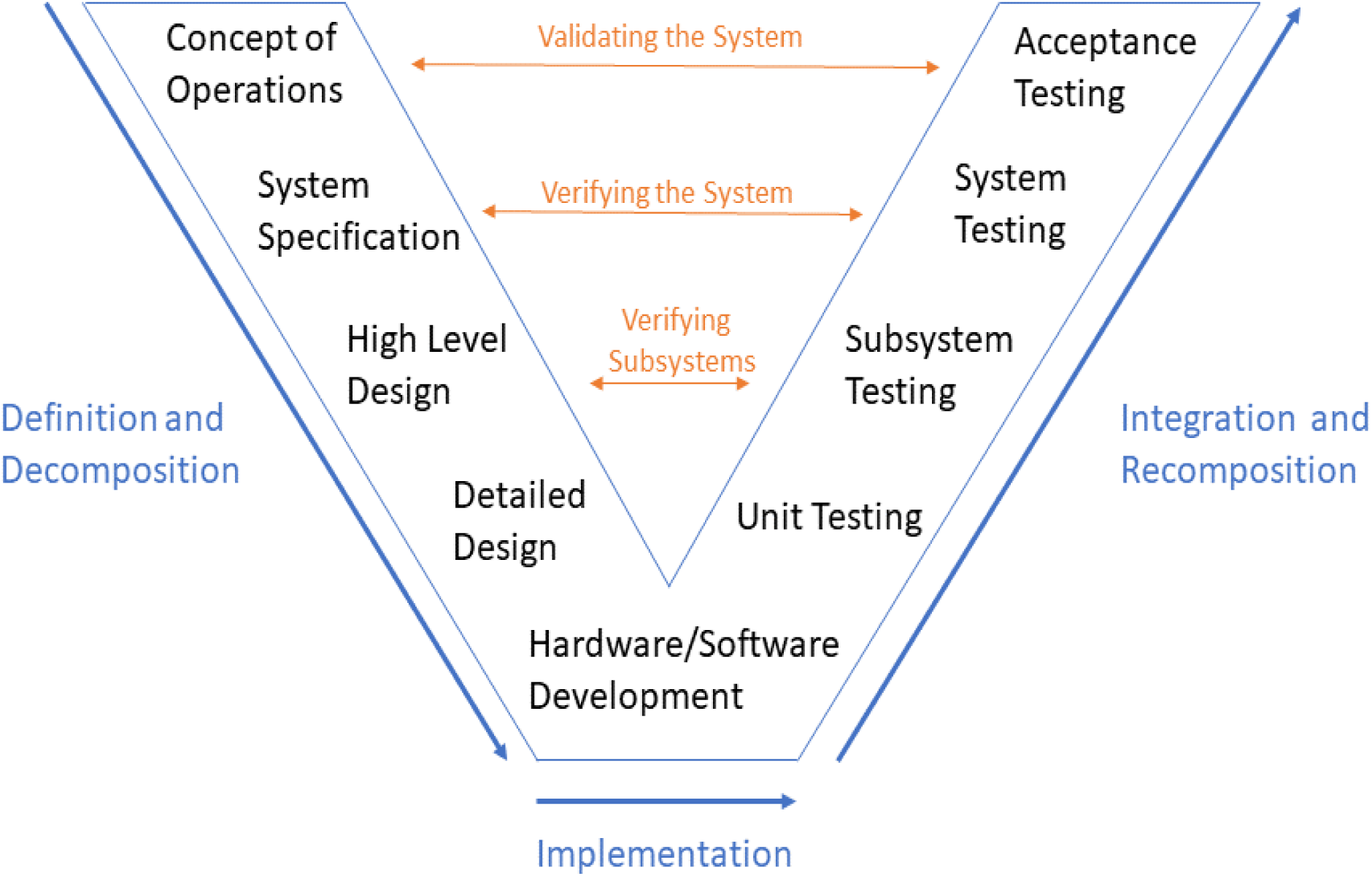
The V-model for the systems engineering process. (Cathleen Shamieh, Systems Engineering for Dummies, Wiley Publishing, Inc., Hoboken, NJ, 2011)

The Design SOPs and Configuration Management system will be applied to the system designed to resolve each specific aim in this project. Each specific aim will begin with a Needs Analysis to determine what the new system needs to be able to do. A requirements analysis will also be conducted to determine what is required to fill those needs. A System Requirements Review (SRR) will demonstrate understanding of the requirements documents (scope, specifications, schedule, validation plans, and budget). SRR will determine the initial design direction and describe preliminary data and progress, and how these will converge to an optimum and complete system configuration for the specific aim. The memory needed to run QB on large datasets and evaluate the performance of the algorithm must be quantified. (Very large sample datasets to test the memory usage and reproducibility of the results of the QB algorithm are being created, as well as to set ranges of parameters for its usage. Parameters include the number of bootstrap replications desired (b), the number of variables in the multivariate analysis (d), and the size of the dataset used (n). Preliminary data indicate that the typical laptop computer can process over one million bootstrap replication samples, independent variables, or sample data points when the other two parameters are minimized, or over one-thousand bootstrap replications, variables, and data points [all maximized at approximately 1000 inputs]).

As the system evolves through the development process, topic experts will be invited to later design reviews (especially CDR, TRR, and MRR). System Design Review (SDR) acts as a control gate that reviews and approves the top-level system design solution and rationale ^19^. It is the decision point to proceed with system specification flow down to individual physical and process configuration items ^19^. System limitations will be refined at SDR.

Using the run-time and memory usage data determined in Specific Aim 1, the performance capabilities of the algorithm on an NSF supercomputer (XSEDE Comet) will be calculated. QB is capable of tackling immensely large datasets, and the computing capabilities of the algorithm will be stretched on a massively parallel machine to demonstrate proof-of-concept. An SDR report will be added to the Design History file for FDA.

A Preliminary Design Review (PDR) will be performed on each configuration item or group of configuration items to: (1) Evaluate the progress, technical acceptability, and risk resolution, (2) Measure its harmony with performance and engineering specialty requirements of the Configuration Item development specification, (3) Evaluate the extent of definition and evaluate the technical risk connected with the selected methods/processes, and (4) Demonstrate the existence and compatibility of the physical and functional interfaces among the configuration item and other items of equipment, facilities, computer software, and personnel ^19^. Topic experts will also be invited to the review. A Blue team and a Red team are used for design and validation, respectively. A PDR report will be added to the Design History file for FDA in the annual reporting system.

Critical Design Review (CDR) is the last design review conducted before an action is taken that is irreversible. (1) CDR is a review to establish that detail design of the configuration item under review meets cost, schedule, and performance requirements. (2) CDR will establish detail design compatibility among the configuration item and other items of equipment, facilities, computer software and personnel. (3) CDR will gauge configuration item risk areas (on a technical performance, cost, and schedule basis). Topic experts will again be invited to the review. A Blue team and a Red team are used for design and validation, respectively. A CDR report will be added to the Design History file for FDA in the annual reporting system.

Deployment Readiness Review (DRR) is held to confirm readiness for deployment. This review is conducted to ensure that all deficiencies are corrected before actual use. The complete system is challenged every feasible way (conceptually, physically, cyber-, etc.). The DRR demands the review and analysis of all subsystem/unit level testing preceding the formal acceptance tests. Topic experts will be invited to the review. A Blue team and a Red team are used for design and validation, respectively. A DRR report will be added to the Design History file for FDA.

The QB algorithm will be translated into Python to make it more accessible to the scientific community. Once QB is available in Python it will run on Amazon Web Services, Microsoft Azure, and Google Compute Engine as well as the NSF XSEDE Comet supercomputer currently being used. The REPLICA algorithm has already been translated from Matlab into Python to make it more accessible to the scientific community. However, the algorithm on which QB relies has yet to be translated and the entirety of the program remains to be validated.

## CREDIT

The preliminary data and proposal described were supported in part by the National Center for Research Resources and the National Center for Advancing Translational Sciences, National Institutes of Health, through Grant UL1TR001998. The content is solely the responsibility of the authors and does not necessarily represent the official views of the NIH. This project was also supported by NSF ACI-1053575 allocation number BIO170011.

## APPENDIX

### REPLICA

~~~
function [BTRAIN,CNTER]=replica(TNSPEC,B)
% TNSPEC=training spectra, B=number of replicates desired.
% REPLICA outputs BTRAIN replicates, and the center of the replicates in CNTER
% “Copyright 2003 Robert A. Lodder & Bin Dai”
[N,D]=size(TNSPEC);
BTRAIN=zeros(B,D);
CNTER=zeros(D,1);
BSAMP=zeros(N,D);
PICKS=rand(B*N,1);
index=find(PICKS==1);
PICKS(index)=0.9999;
PICKS=reshape(PICKS,B,N);
PICKS=fix(N*PICKS+1);
for I=1:B
   BSAMP=TNSPEC(PICKS(I,:),:);
   BTRAIN(I,:)=sum(BSAMP)/N;
end
BTRAIN;
CNTER=sum(BTRAIN)/B;
~~~

### SOB

~~~
function [ECDF,TCDF,R]=SOB(BTRAIN,CTN,NTN,TESTSPEC)
% This routine takes 2 spectral groups (with = numbers of samples)
% and calculates an ECDF and TCDF for QQ plotting. The routine
% also returns a correlation coefficient between the 2 CDFs.
% Training replicates and center must be provided, along with
% test spectra. BNUM = the number of replicates used.
% Note: you may have to set EPSILON(1e-200) or greater to prevent
% underflow errors in high dimensional hyperspaces
% Copyright 2003 Dr. Robert.A.Lodder & Bin Dai
[NTEST,C]=size(TESTSPEC);
[BTEST,CTEST]=replica(TESTSPEC,NTN);
[M,C]=size(BTRAIN);
p=0.1;
CTNN=repmat(CTN,M,1);
ST=(sum(((BTRAIN-CTNN).^C)′)).^(1/C);
CTNN=repmat(CTN,NTN,1);
SX=(sum(((BTEST-CTNN).^C)′)).^(1/C);
M=2*M;
TRIM=fix((M*p)+1):fix(M-(M*p));
ECDF=[ST,SX];
TCDF=[ST,ST];
ECDF=sort(ECDF);
TCDF=sort(TCDF);
ECDF=ECDF(TRIM);
TCDF=TCDF(TRIM);
R=corrcoef(TCDF,ECDF)
~~~

